# Adolescent Social Isolation Facilitates Tau Spread in Raphe Nuclei, Linking Depression and Hyperalgesia in Alzheimer’s Disease

**DOI:** 10.64898/2026.01.23.701108

**Authors:** Nagalakshmi Balasubramanian, Gabriel Gaudencio, Kanza M. Khan, Natalie Biggerstaff, Catherine A Marcinkiewcz

## Abstract

Tau pathology in brainstem serotonergic circuits drives early neuropsychiatric dysfunction in Alzheimer’s disease (AD), yet mechanisms linking the exposome, particularly social stress exposures to depression and altered pain perception, remain unclear. Here, we demonstrate that adolescent social isolation, a critical psychosocial exposome factor and major trigger of depression, facilitates tau propagation in the dorsal raphe nucleus (DRN) and downstream raphe nuclei, producing both neuropsychiatric and pain-related sequelae.

Tau[P301L] was transduced into the DRN of 30-day socially isolated or group housed C57BL/6J mice using AAV, with control mice receiving AAV-GFP. Four weeks post-transduction, anxiety, social behavior, and pain sensitivity were assessed, and phosphorylated tau (ptau) within the DRN and spread to the median raphe (MRN) and raphe magnus (RMg) serotonergic neurons were evaluated. Socially isolated tau[P301L] mice exhibited hyperlocomotion, anxiety-like behavior, social deficits and hyperalgesia. Histological analysis revealed elevated ptau within TPH2 neurons in the DRN and trans-synaptic tau spread to MRN and RMg, accompanied by reduced TPH2 and increased ptau at the downstream raphe nuclei. Fluorescence *in situ* hybridization confirmed altered expression of *Slc6a4* and *Tgm2*, a stress-responsive gene implicated in AD. These results indicate that DRN tau under social stress drives neuropsychiatric phenotypes, while tau spread to MRN and RMg disrupts serotonergic modulation of pain. This study provides the first evidence that adolescent stress promotes tau propagation within the raphe nuclei, linking early neuropsychiatric and pain-processing deficits to prodromal Alzheimer’s disease and identifying a critical pathway through which psychosocial exposome risk converges with tau pathology to enable early intervention.

## Introduction

Alzheimer’s disease (AD) is a devastating neurodegenerative disorder characterized by progressive cognitive decline. Yet long before cognitive symptoms emerge, many individuals experience profound neuropsychiatric and sensory disturbances, including depression, anxiety, social withdrawal, and altered pain sensitivity [1–3]. These early symptoms not only diminish quality of life but may also actively contribute to disease progression. Chronic pain is increasingly recognized as a clinically significant comorbidity in early AD. Epidemiological data indicates that pain is reported by 40.4% of nursing home residents (NHRs) without cognitive impairment and by 34% of those with mild cognitive impairment, compared with only 24.9% of residents with severe cognitive impairment [4,5], suggesting an apparent decline in pain perception or reporting with increasing cognitive impairment. Intriguingly, AD and chronic pain share overlapping risk factors such as aging, family history, depression, and social isolation.

Within the broader framework of exposome, social isolation or loneliness during adolescence represents a potent psychosocial stressor capable of altering neurodevelopmental trajectories and increasing susceptibility to both mood disorders and neurodegenerative disease [6].

However, how adolescent stress exposures mechanistically interface with AD pathology remains insufficiently defined.

Tau aggregation and accumulation is a hallmark of AD neuropathology and is strongly correlated with neurodegenerative progression. While tau pathology has traditionally been associated with cortical and hippocampal degeneration, recent human postmortem studies reveal early tau deposition within brainstem serotonergic nuclei, particularly the dorsal raphe nucleus (DRN) [7–9]. DRN is a major source of forebrain serotonin and is essential for regulating mood, social behavior, pain modulation, and stress adaptation [10]. Disruption of serotonergic circuits has been implicated in depression, anxiety, and altered nociception, all of which commonly emerge in the prodromal stages of AD [11]. Studies including our own report demonstrate phosphorylated tau (ptau) deposition in the DRN of cognitively normal individuals, with greater accumulation observed in AD individuals, indicating that tau pathology initiates in serotonergic circuits prior to overt cognitive decline [7–9,12,13]. Furthermore, our experimental studies using mice expressing human wild-type MAPT or AAV-mediated tau[P301L] selectively within the DRN revealed robust social deficits in the absence of cognitive impairment [9,13].

These findings suggest that early tau driven serotonergic dysfunction may render the brain more vulnerable to environmental stressors. Furthermore, trans-synaptic propagation of tau to other brain areas [14–16] is enhanced by neural activity and stress [17,18]. Given the extensive reciprocal connectivity among raphe nuclei, tau pathology from the DRN may propagate trans-synaptically to the median raphe nucleus (MRN) and raphe magnus (RMg) which are serotonergic hubs that contribute to mood regulation and descending pain control pathways.

Consequently, stress-enhanced tau spread across these circuits could drive a spectrum of neuropsychiatric and pain-related phenotypes seen in AD. Yet this hypothesis has not been rigorously tested.

In the present study, we tested the hypothesis that adolescent social isolation, as a key exposome factor, facilitates tau propagation within raphe circuits, linking early-life stress to neuropsychiatric and pain-related consequences in AD. We used an AAV-based strategy to express tau[P301L] in the DRN of socially isolated C57BL/6J mice during adolescence. We assessed anxiety-like behavior, sociability, thermal and mechanical nociception, and quantified tau burden in the DRN and trans-synaptic spread to the MRN and RMg. Through combined behavioral, molecular, and histochemical analyses, we demonstrate that adolescent isolation stress exposure amplifies tau spread across brainstem serotonergic networks and produces persistent affective and pain-related phenotypes. These findings establish a mechanistic bridge between exposome and early AD pathogenesis. By revealing how psychosocial stress interacts with tau-driven dysfunction in serotonergic circuits, this work highlights a critical window where environmental risk factors shape long-term vulnerability, offering opportunities for prevention and early therapeutic intervention.

## Methods

### Ethical approval

All animal procedures were reviewed and approved by the University of Iowa (UI) Office of Animal Resources and University of Florida that abided by the AVMA and NIH.

### Animals

Male C57BL/6J mice (Jackson Labs Stock 000664) were purchased for arrival at PND 21, shipped five mice per cage. At age PND 25, half of each cohort (i.e., 10 mice) was moved to individual housing. A total of 40 C57BL/6J male mice were used in the experiment. Mice were housed in a temperature- and humidity-controlled, AALAC-approved vivarium at the University of Iowa on a standard 12h/12h dark/light (reverse) cycle and in accordance with institutional requirements for animal care. Mice were individually or group housed in conventional-style cob bedding rodent cages with nestlets containing separate food and water that could be obtained *ad libitum*.

### Stereotaxic surgery

Surgeries were performed as previously described [9]. Briefly, at P56 (i.e after five weeks of isolation), mice were subjected to intracranial surgery for delivering AAV carrying mutant tau[p301l] or eGFP (Fig. 1A). Briefly the mice were induced into deep anesthesia using 3% (v/v) isoflurane in oxygen and placed in a Leica Angle 2 stereotaxic apparatus (Germany) on a temperature-controlled heating pad. Anesthesia was maintained at a concentration of 1.5-2% isoflurane throughout the procedure. AAV8-CBA-P301L-tau-WPRE or AAV8-CAG-eGFP-WPRE (control) vectors were precisely injected into the DRN (coordinates: ML: 0.0; AP: -4.65; DV: -3.30 relative to Bregma, with a 23.58° tilt from the ML plane) using a 1 µL Hamilton syringe at an infusion rate of 100 nL/min, for a total volume of 500 nL. Post-operative pain management was provided by administering two meloxicam injections (0.4 mg/kg, subcutaneous or intraperitoneal) 24 hours apart. Mice were carefully monitored following surgery and returned to their pre-surgical housing conditions.

**Figure 1.**
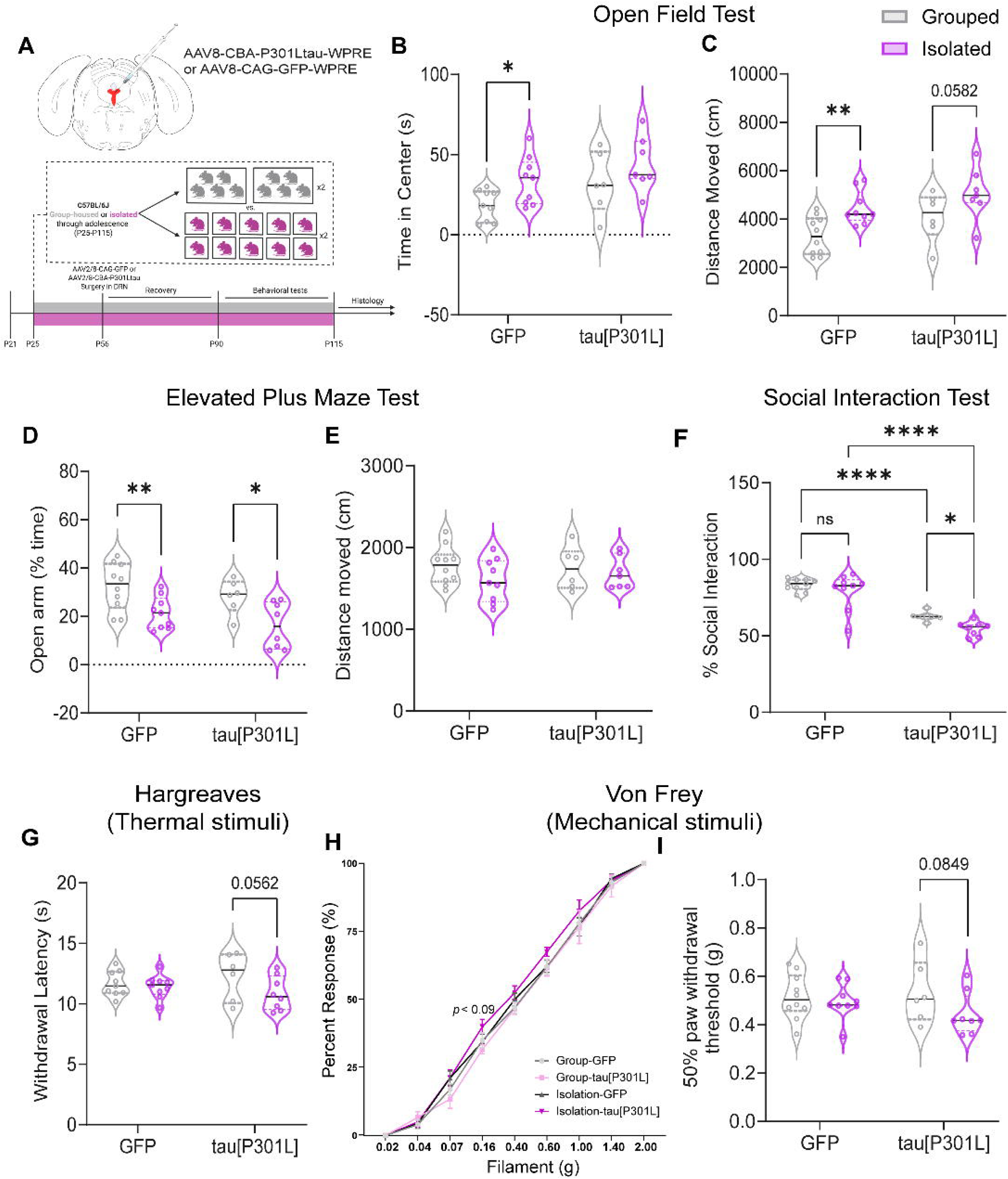
Adolescent social isolation and the impact of tau[P301L] expression on anxiety, social behavior, and pain sensitivity. (A) Experimental timeline illustrating the period of adolescent social isolation, intracranial infusion of AAV8-CBA-P301Ltau-WPRE or AAV8-CAG-GFP-WPRE into the dorsal raphe nucleus (DRN), and subsequent behavioral testing schedule for anxiety, depression, and pain related measures. (B-C) Open field test showing (B) time spent in the center and (C) total distance moved in group-housed and socially isolated GFP and tau[P301L] mice. (D-E) Elevated plus maze test showing (D) percentage of time spent in the open arms and (E) total distance moved across groups. (F) Percentage of interaction time with a stranger mouse in the three-chamber social interaction test for group-housed and socially isolated GFP and tau[P301L] mice. (G-I) Pain sensitivity assessed using (G) the Hargreaves test showing paw withdrawal latency to thermal stimuli, and (H-I) the von Frey test showing percent response and 50% paw withdrawal threshold to mechanical stimuli in group-housed and socially isolated GFP and tau[P301L] mice. Values (n =7-10/group) are represented as means (±SEM), and the data were analyzed by two-way ANOVA (\**p* < 0.05, \*\**p* < 0.01, \*\*\*\**p* < 0.0001). Part of the figure was made in BioRender.

### Viral constructs

Viral constructs were prepared as previously described [9]. The plasmid utilized to create the P301L-tau viral vector was kindly donated by Gloria Lee, while the AAV2/8-CBA-P301Ltau-WPRE construct was prepared by the Viral Vector Core Facility at the University of Iowa. The AAV2/8-CAG-GFP-WPRE control virus was sourced from the same facility.

### Behavior Assays

All behavioral testing was conducted in the span of four weeks following surgery, and the assays were performed in the following sequence: Open Field test, Elevated Plus Maze test, Social Interaction test, Von Frey test, and Hargreaves test. At least two-day gap was allowed between each behavioral test. All the behavior tests were conducted between 8 AM-12 PM. The experimenters were blinded to the group assignments of the animals during data collection and analysis until statistical comparisons were made.

#### Elevated Plus Maze

The Elevated Plus Maze (EPM) test was performed as previously described [9,19]. Briefly, mice were placed in the center of an elevated, plus-shaped maze consisting of two open arms, two enclosed arms (5 x 35 cm), and a neutral starting zone (5 x 5 cm), positioned 60 cm above the floor. The enclosed arms had 20 cm high walls, providing a dark, sheltered space for the animals. The maze was illuminated with overhead LEDs, maintaining ∼20 lux in the open arms and < 5 lux in the closed arms. Mice were allowed to explore the maze freely for 5 mins, with the experimenter obscured from view by an opaque curtain. Sessions were recorded using a top view Basler GenICam camera and Media Recorder 4 software (Noldus, VA, USA). Distance traveled, and the % time spent in the open and closed arms, were analyzed using EthoVision XT15 (Noldus).

#### Open Field Test

The Open Field test (OFT) was performed as previously described [9]. Briefly, mice were placed in the corner of a white open box (L x W x H: 50 x 50 x 25 cm), situated in a custom-made sound-attenuating chamber. The arena was illuminated with dim lighting (∼20 lux in the center of the arena), and behavior was recorded with a camera mounted above. The first 10 minutes of each session were analyzed to assess parameters. The total distance traveled (cm) and time spent in the center of the arena were recorded and analyzed with Ethovision XT14 software.

#### Social Interaction

The Social Interaction (SI) test was performed as previously described [9]. Briefly, the experimental mouse was placed in the central chamber of a transparent 3-chamber arena (illuminated to 20 lux) and allowed to explore the environment for 10 minutes. Then, a novel C57BL/6J mouse of the same sex and age (stranger mouse) was introduced into one of the side chambers inside a metal holding cage, while an empty metal cage was placed in the opposite chamber. The experimental mouse was given 10 minutes to explore the environment and interact with both the stranger mouse and the empty cage. To control potential side preferences, the position of the stranger mouse was alternated between the left and right chambers. The total time spent interacting with the stranger mouse and the empty cage was recorded.

#### Von Frey

The von Frey test was performed as previously described [20,21]. Briefly, mice were placed in a clear acrylic box (10 x 10 x 15 cm) and allowed to acclimate to the test environment, which included a raised wire mesh platform with 5 x 5 mm holes, for at least 3 hours on two consecutive days. Mechanical sensitivity was assessed by applying von Frey filaments (Stoelting, Wood Dale, IL, USA) of varying forces to the plantar surface of each hind paw. The number of responses to each filament, out of five applications, was recorded and used to calculate the 50% withdrawal threshold for each paw and percent response.

#### Hargreaves Test

The Hargreaves test was performed as previously described by us [20,21]. Mice were allowed to acclimate in a clear acrylic box (10 x 10 x 15 cm) on the test apparatus (IITC Life Science heated base, Model 400), with the temperature maintained at 30°C, for at least three hours on two consecutive days prior to testing. Thermal sensitivity was assessed by directing a heat-generating light beam onto the plantar surface of each hind paw. The time taken to elicit a paw withdrawal response was measured three times per paw, and the average latency was calculated for each paw.

## Histology

Mice were anesthetized with tribromoethanol (250 mg/kg, i.p.) and transcardially perfused with 30 mL of 0.01 M phosphate-buffered saline (PBS), followed by 30 mL of 4% paraformaldehyde (PFA) after 8 weeks of social isolation. Brains were then extracted and post-fixed for 24 hours in 4% PFA at 4°C. After post-fixation, brains were transferred sequentially to 15% and 30% sucrose solutions and allowed to equilibrate. The brains were then embedded in OCT and sectioned at 45 μm using a frozen sliding microtome (Leica Biosystems).

## Immunofluorescent staining

Immunofluorescence was performed to measure TPH2 and phosphorylated tau (pSer202/Thr205) levels in the serotonin neurons within DRN as described previously [13]. Briefly, for each region of interest (ROI) such as DRN, MRN and RMg, 4-6 slices were used across the rostral-caudal axis. Slices were washed in PBS 3 times for 5 minutes, then permeabilized using 0.5% Triton X-100/PBS for 30 min, blocked in 10% normal donkey serum in 0.1% Triton X-100/PBS. Tissue was rinsed 2 times for 5 minutes and then incubated with the respective primary and secondary antibodies (Table 1). Slices were subsequently washed in PBS 4 times for 10 minutes, mounted on glass slides, and coverslipped with Vectashield mounting media (Vector Laboratories, Inc.). Images were acquired on an Olympus FV-3000 confocal microscope with z-stacks (0.5 μm/step) at 20X or 40X with an optical resolution factor of 1X, 2X, or 3X. Multiple tiles were imaged for every region of interest, and mosaic stitching was performed using FV3000 Multi-Area Time-Lapse (MATL) Software. After imaging, confocal stacks were converted to maximum projection images using Image J software. Orthogonal images were also generated in the Image J software to demonstrate the accumulation of ptau in the serotonin neurons.

**Table 1.** Antibodies used for immunofluorescence.

| Target | Reagents |
| --- | --- |
| Tau<br>(pSer202/Thr205,<br>AH36) | Primary: Rabbit anti-tau, StressMarq SMC-601, lot MK287130, 1:2000.<br>Secondary: Alexa 555 donkey anti-rabbit, Invitrogen A31572, 1:1000. |
| TPH2 | Primary: Goat anti-TPH2, ARP Eb11012, lot P1, 1:500.<br>Secondary: Alexa 647 donkey anti-goat, Jackson 705-605-003, 1:800. |

## Image Processing and Analysis

Maximum projection confocal images from Image J were saved as TIFF files before being imported into QuPath software (Version 0.5.1). In QuPath, all images were preprocessed by adjusting the brightness and contrast to consistent levels. Analysis was performed with the experimenter blinded to group assignment.

### TPH2 cell density

For object detection, an object classifier was set of TPH2 expressing neurons. The TPH2 cell density was determined by calculating the number of TPH2 positive cells per unit area (cells/μm²) within the DRN, MRN, and RMg. The total count of TPH2 cells was divided by the corresponding area of the ROI in each brain slice, and an average density value was obtained for each brain.

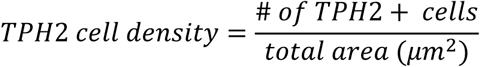

### TPH2 % Immunoreactivity (IR)

A pixel classifier was set for TPH2 fluorescence signals and TPH2 % IR was calculated as the proportion of the area occupied by TPH2 immunoreactive signal (Tph2 area) relative to the total area (DRN total area). This measure quantifies the overall signal of TPH2 within the region, independent of individual cell counts. The same procedure was applied to the MRN and RMg.

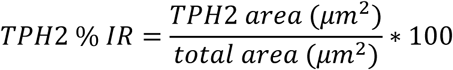

### AH36 % Immunoreactivity (IR)

A pixel classifier was set for AH36 (ptau) fluorescence signals and AH36 % IR was calculated as the proportion of the area occupied by AH36 immunoreactive signal (AH36 area) relative to the total area (DRN total area). This measure quantifies the overall expression of AH36 staining within the region, independent of individual cell counts. The same procedure was applied to the MRN and RMg.

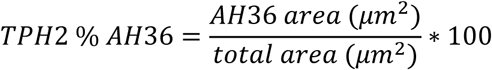

### AH36 % IR in TPH2+ neurons

The percentage of AH36 immunoreactivity (% IR) within TPH2+ neurons was measured using pixel classifier. First, a TPH2 pixel classifier was created to segment and quantify the total TPH2+ area (μm²) in the DRN, MRN, and RM. This area was then used as an annotation to define the serotonergic neuron region of interest. A second AH36 pixel classifier was applied within the TPH2 defined area to detect AH36+ (phosphorylated tau) puncta that exceeded the set threshold. The % IR AH36 within TPH2 neurons was calculated as the area of AH36 signal inside the TPH2+ region divided by the total TPH2+ area, multiplied by 100.

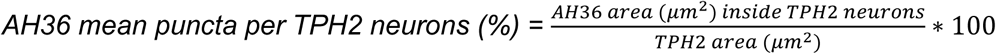

## RNAscope fluorescent in situ hybridization (FISH)

FISH was performed using the RNAscope Multiplex Fluorescent v2 assay (Advanced Cell Diagnostics) according to the manufacturer’s guidelines, with minor modifications. Coronal brain sections (45 µm) were mounted on Histobond Plus slides (VWR), air dried, and baked at 60°C for extended time of 90 minutes. Slides were post-fixed in pre-chilled 10% neutral buffered formalin (NBF) for 60 minutes at 4°C, dehydrated through an ethanol gradient (50%, 70%, 100%), and baked again at 60°C for 60 minutes. Sections were treated with RNAscope Hydrogen Peroxide for 10 minutes at room temperature, followed by heat-mediated target retrieval in 1X Target Retrieval Reagent for 10 minutes. Slides were then washed in 100% ethanol, air dried, and a hydrophobic barrier was drawn around tissue sections. Protease III was applied for 45 minutes at 40°C in a HybEZ oven. After the pretreatment steps, respective mRNA probes for *Tgm2*, *Slc6a4*, and *Tph2* (Table 2) were hybridized at 40°C for 3 hours, followed by sequential amplification (AMP1-AMP3) and horseradish peroxidase (HRP) channel development. Fluorescent detection was achieved using Opal dyes (Table 2), with HRP-blocking steps between each fluorophore deposition. Sections were counterstained with DAPI, coverslipped with Vibrance mounting medium (Vector Laboratories), and stored in the dark until imaging. The slides were imaged using an Olympus SLIDEVIEW VS200 digital slide scanner and the images were analyzed in QuPath (version 0.5.1). *Slc6a4* and *Tgm2* expression within TPH2 neurons were measured by creating a pixel classifier first for TPH2 and used this as a parent annotation for measuring the puncta of *Slc6a4* and *Tgm2* using their respective pixel classifier.

**Table 2.** RNAscope probes and Opal dyes

| Target gene | Reagents |
| --- | --- |
| *Tgm2* | Probe: Mm-Tgm2-C1 (ACDbio, cat# 537801), dilution 1:50.<br>Fluorophore: Opal 650 (Akoya, cat# OP-001005), dilution 1:1000. |
| *Slc6a4* | Probe: Mm-Slc6a4-C3 (ACDbio, cat# 315851-C3), dilution 1:50.<br>Fluorophore: Opal 570 (Akoya, cat# OP-001003), dilution 1:1000. |
| *Tph2* | Probe: Mm-Tph2-C4 (ACDbio, cat# 318691-C4), dilution 1:100.<br>Fluorophore: Opal 520 (Akoya, cat# OP-001001), dilution 1:2000. |

## Statistical Analysis

Statistical tests were conducted using Prism 9 (GraphPad, La Jolla, CA). All data sets were assessed for normality and homogeneity of variance prior to analysis. A two-way ANOVA was performed to compare cell means across the two independent variables. For each comparison, the means within each row and column were compared across the four family groups (one per row and one per column). No correction for multiple comparisons was applied, and each comparison was evaluated independently. Fisher’s Least Significant Difference (LSD) test was used for post hoc comparisons where applicable. Statistical significance was ascertained when p<0.05. Statistical significance was defined as \**p <* 0.05, \*\**p <* 0.01, \*\*\**p <* 0.001, \*\*\*\**p <* 0.0001. Data are expressed as mean ± SEM unless otherwise specified.

## Results

### Adolescent social isolation and the impact of ptau accumulation within 5HT-DRN neurons in anxiety, depression and nociceptive behavior

Adolescent social isolation is a major early-life stressor that profoundly impacts mental and physical health, contributing to anxiety, depression, and altered nociception later in life.

Serotonergic neurons in the DRN play a critical role in modulating these behaviors through projections to various subcortical regions, including other raphe nuclei such as the median raphe nucleus (MRN) and raphe magnus (RMg), which have been implicated in tau accumulation during early AD. To examine the interaction between stress and AD-related tau pathology on neuropsychiatric and pain phenotypes, we intracranially injected AAV expressing tau[P301L] or GFP into the DRN of mice that underwent adolescent social isolation or group housing, followed by behavioral assessment for anxiety, depression, and nociception (Fig. 1A).

Analysis of the open field test showed that adolescent social isolation increased both the time spent in the center zone and overall locomotor activity. Two-way ANOVA revealed significant main effects of virus genotype (*F*(1,27) = 4.968, *p* = 0.0343) and housing condition (*F*(1,27) = 6.178, *p* = 0.0194), with no significant interaction (*F*(1,27) = 0.081, *p* = 0.7776) for time spent in the center. Post hoc Fisher’s LSD analysis indicated that isolated GFP mice spent significantly more time in the center than grouped GFP controls (*p* = 0.0413), whereas tau[P301L] mice showed no significant differences between housing conditions (Fig. 1B). Similarly, isolation induced hyperlocomotion in both genotypes. Two-way ANOVA for distance moved revealed significant effects of virus genotype (*F*(1,27) = 5.864, *p* = 0.0224) and housing condition (*F*(1,27) = 12.65, *p* = 0.0014), without interaction (*F*(1,27) = 0.245, *p* = 0.6247). Post hoc analysis confirmed that isolated GFP (*p* = 0.0041) and tau[P301L] mice (*p* = 0.0582) were hyperlocomotive compared to their grouped counterparts (Fig. 1C).

Analysis of the EPM test revealed that adolescent social isolation stress induced anxiety-like behavior in both GFP and tau[P301L] mice. Two-way ANOVA showed a significant main effect of housing condition (*F*(1,29) = 14.65, *p =* 0.0006), but no significant effects of virus genotype (*F*(1,29) = 2.405, *p =* 0.1318) or interaction between the two factors (*F*(1,29) = 0.03548, *p =* 0.8519) on % time spent in the open arms. Post hoc Fisher’s LSD analysis indicated that both isolated GFP (*p* = 0.0088) and tau[P301L] (*p* = 0.0133) mice spent significantly less time in the open arms compared to their respective group-housed controls (Fig. 1D). The distance moved parameter remained unchanged across both isolated and grouped GFP or tau[P301L] mice (Fig. 1E).

Next, we investigated the combined effects of adolescent social isolation and DRN-tau[P301L] expression on social behavior. Previously, we observed pronounced social deficits in both tau[P301L] [9] and htau [13] mice compared with their respective controls. Analysis of the SI test revealed that adolescent social isolation induced social deficits selectively in tau[P301L] mice.

Two-way ANOVA demonstrated a significant main effect of viral genotype (*F*(1,29) = 81.41, *p* < 0.0001) and housing condition (*F*(1,29) = 5.847, *p* = 0.0221), with no significant interaction between these factors (*F*(1,29) = 0.7303, *p* = 0.3998) on the % social interaction time. Post hoc Fisher’s LSD analysis indicated that socially isolated tau[P301L] mice exhibit reduced % social interaction time compared to their group-housed counterparts (*p* = 0.0402). Furthermore, both group-housed and isolated tau[P301L] mice exhibited pronounced social deficits relative to GFP-expressing group-housed (*p* < 0.0001) and isolated controls (*p* < 0.0001), respectively (Fig. 1F).

Pain sensitivity was assessed using the Hargreaves and von Frey tests to measure responses to thermal and mechanical stimuli, respectively. In the Hargreaves test, two-way ANOVA revealed no significant effects of virus genotype (*F*(1,28) = 0.0022, *p* = 0.9629), housing condition (*F*(1,28) = 3.129, *p* = 0.0878), or their interaction (*F*(1,28) = 1.520, *p* = 0.2279) on paw withdrawal thresholds. However, the post hoc analysis showed an increasing trend in thermal sensitivity in isolated tau[P301L] mice compared to grouped mice (*p* = 0.0562) (Fig. 1G), while no significant differences were seen between grouped and isolated mice within GFP mice (*p* = 0.6867). In the Von Frey test, two-way ANOVA for V50 paw averages showed no significant effect of virus genotype (*F*(1,29) = 0.2742, *p* = 0.6045) and housing (*F*(1,29) = 2.722, *p* = 0.1098) (Fig. 1H). Post hoc comparisons revealed no change in the isolated GFP mice, while isolated tau[P301L] mice exhibit lower V50 paw withdrawal threshold value as compared to control group mice (*p* = 0.0849), suggesting isolated tau[P301L] mice are sensitive to mechanical stimuli (Fig. 1I).

### Tau pathology in the dorsal raphe nucleus is exacerbated by adolescent social isolation

Next, we examined the effects of adolescent social isolation on TPH2 neuron density and ptau (AH36) accumulation within the DRN (Fig. 2A-G). Analysis of TPH2 immunoreactivity revealed that isolated tau[P301L] mice exhibited higher TPH2-positive area compared to isolated GFP controls, with an increasing trend relative to group-housed tau[P301L] mice. Two-way ANOVA revealed a significant main effect of virus genotype (*F*(1,29) = 7.186, *p* = 0.0120), but no main effect of housing (*F*(1,29) = 1.556, *p* = 0.222) or interaction between housing x virus genotype (*F*(1,29) = 1.887, *p* = 0.780). Post hoc Fisher’s LSD analysis confirmed that isolated tau[P301L] mice displayed significantly higher TPH2 immunoreactive area than isolated GFP mice (*p* = 0.0061), with a trend toward increased levels compared to group-housed tau[P301L] mice (*p* = 0.0959) (Fig. 2D). Similarly, TPH2+ cell density in the DRN was elevated in isolated tau[P301L] mice relative to both isolated GFP and group-housed tau[P301L] mice (Fig. 2E). Two-way ANOVA revealed a significant interaction between housing x virus genotype (*F(*1,29) = 4.559, *p* = 0.0413). Post hoc analysis showed that isolated tau[P301L] mice had significantly higher TPH2 cell density compared to isolated GFP mice (*p* = 0.0088) and group-housed tau[P301L] mice (*p* = 0.0182). We next assessed ptau (AH36) accumulation within the DRN. Overall AH36-immunoreactive area was elevated in tau-injected mice regardless of housing condition (Fig. 2F). However, when ptau levels were specifically analyzed within TPH2+ neurons, a pronounced increase was observed in isolated tau[P301L] mice compared to all other groups. Two-way ANOVA revealed significant main effects of virus genotype (*F*(1,29) = 33.60, *p* < 0.0001) and housing condition (*F*(1,29) = 8.731, *p* = 0.0062), as well as a significant interaction between housing x virus genotype (*F*(1,29) = 9.421, *p* = 0.046). Post hoc Fisher’s LSD analysis confirmed that isolated tau[P301L] mice displayed markedly higher ptau levels within TPH2 neurons compared to group-housed tau[P301L] mice (*p* = 0.0005) (Fig. 2G).

**Figure 2.**
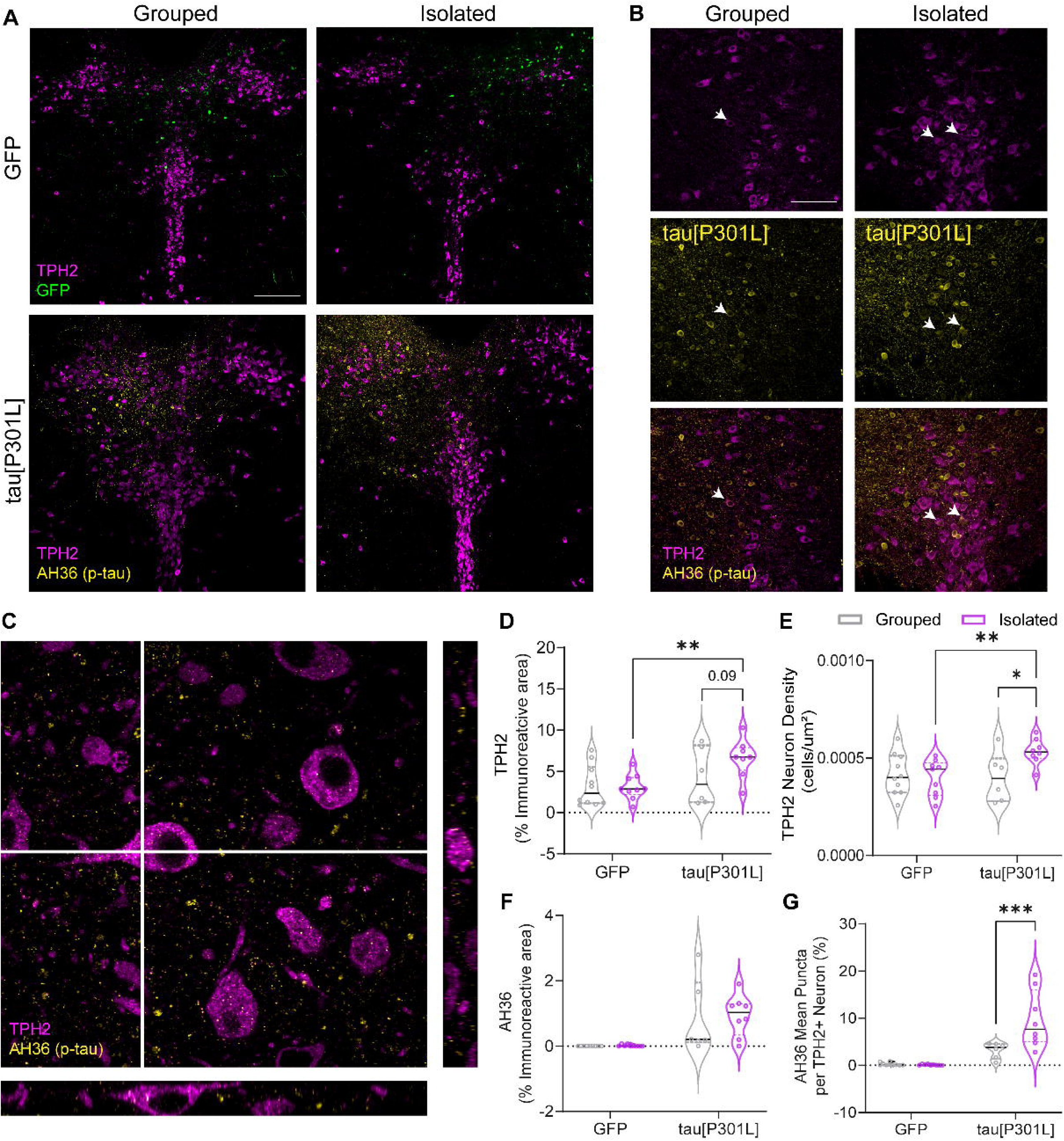
Adolescent isolation stress increases tau burden in the dorsal raphe nucleus (DRN) and enhances ptau accumulation within serotonergic neurons. (A) Representative confocal images showing increased AH36 (ptau, pSer202/pThr205) immunofluorescence in the DRN of adolescent socially isolated tau[P301L] mice compared to group-housed and GFP controls. Scale bar = 200 µm. The top two panels show GFP control: group-housed and isolated conditions, while the bottom two panels show tau[P301L]-group-housed and isolated conditions. (B) Inset images showing ptau accumulation within 5-HT neurons in the DRN: top, TPH2 (serotonergic marker); middle, AH36 (ptau, pSer202/pThr205); bottom, merged overlay. Scale bar = 50 µm. (C) Orthogonal projection image depicting ptau accumulation within DRN TPH2-positive neurons. (D-G) Quantification showing the effects of adolescent social isolation on (D) percentage of TPH2-immunoreactive area, (E) TPH2 neuron density, (F) percentage of AH36-immunoreactive area, and (G) mean number of AH36 (ptau) puncta per TPH2-positive neuron in GFP and tau[P301L] mice. Values (n =7-10/group) are represented as means (±SEM), and the data were analyzed by two-way ANOVA (\**p* < 0.05, \*\**p* < 0.01, \*\*\**p* < 0.001).

We then analyzed expression levels of *Slc6a4* and *Tgm2*, which encode the serotonin transporter (SERT) and transglutaminase 2 (a stress-related protein known to induce tau aggregation), to mechanistically link adolescent social isolation and tau pathology-induced increases in TPH2 levels within the DRN (Fig. 3A-E). We found that stress selectively upregulated *Slc6a4* expression in tau[P301L] mice, accompanied by a concomitant increase in *Tph2* mRNA levels (Fig. 3A-C). Two-way ANOVA for *Tph2* gene expression revealed a significant main effect of virus genotype (*F(*1,19) = 5.942, *p =* 0.0248), no main effect of housing condition (*F(*1,29) = 8.731, *p =* 0.0062), and no significant interaction between housing x virus genotype. Post hoc Fisher’s LSD analysis indicated significantly higher *Tph2* mRNA levels in isolated tau[P301L] mice compared to isolated GFP mice (*p =* 0.0315) (Fig. 3B). Similarly, two-way ANOVA for *Slc6a4* gene expression revealed a significant main effect of housing condition (*F(*1,19) = 12.48, *p =* 0.0222), with no significant effects of virus genotype or housing x virus interaction. Post hoc Fisher’s LSD analysis showed significantly higher *Slc6a4* mRNA levels in isolated tau[P301L] mice compared to group-housed tau[P301L] mice (*p =* 0.030), and a trend toward an increase relative to isolated GFP mice (*p =* 0.069) (Fig. 3C). *Tgm2* is widely known for its role in stress and AD [22–25], and we have previously reported elevated *TGM2* levels in DRN of another AD model, *Htau*, which expresses human wild-type MAPT [13]. In the present study, we found that social isolation stress increased *Tgm2* mRNA expression in both GFP and tau[P301L] mice (Fig. 3D-E). Two-way ANOVA for *Tgm2* gene expression revealed a significant main effect of housing condition (*F(*1,19) = 16.99, *p =* 0.0006), with no effects of virus genotype or housing x virus genotype interaction. Post hoc Fisher’s LSD analysis confirmed significantly higher *Tgm2* mRNA levels in both isolated GFP (*p =* 0.0150) and tau[P301L] mice (*p =* 0.0051) compared to their respective group-housed counterparts (Fig. 3D-E). These results suggest that *Tgm2* is relatively insensitive to the phosphorylation state of tau but is highly responsive to stress. Thus, by regulating transcription of *Tgm2* and determining its relative abundance in the cell, the exposome may be able to convert resilient neurons into vulnerable neurons.

**Figure 3.**
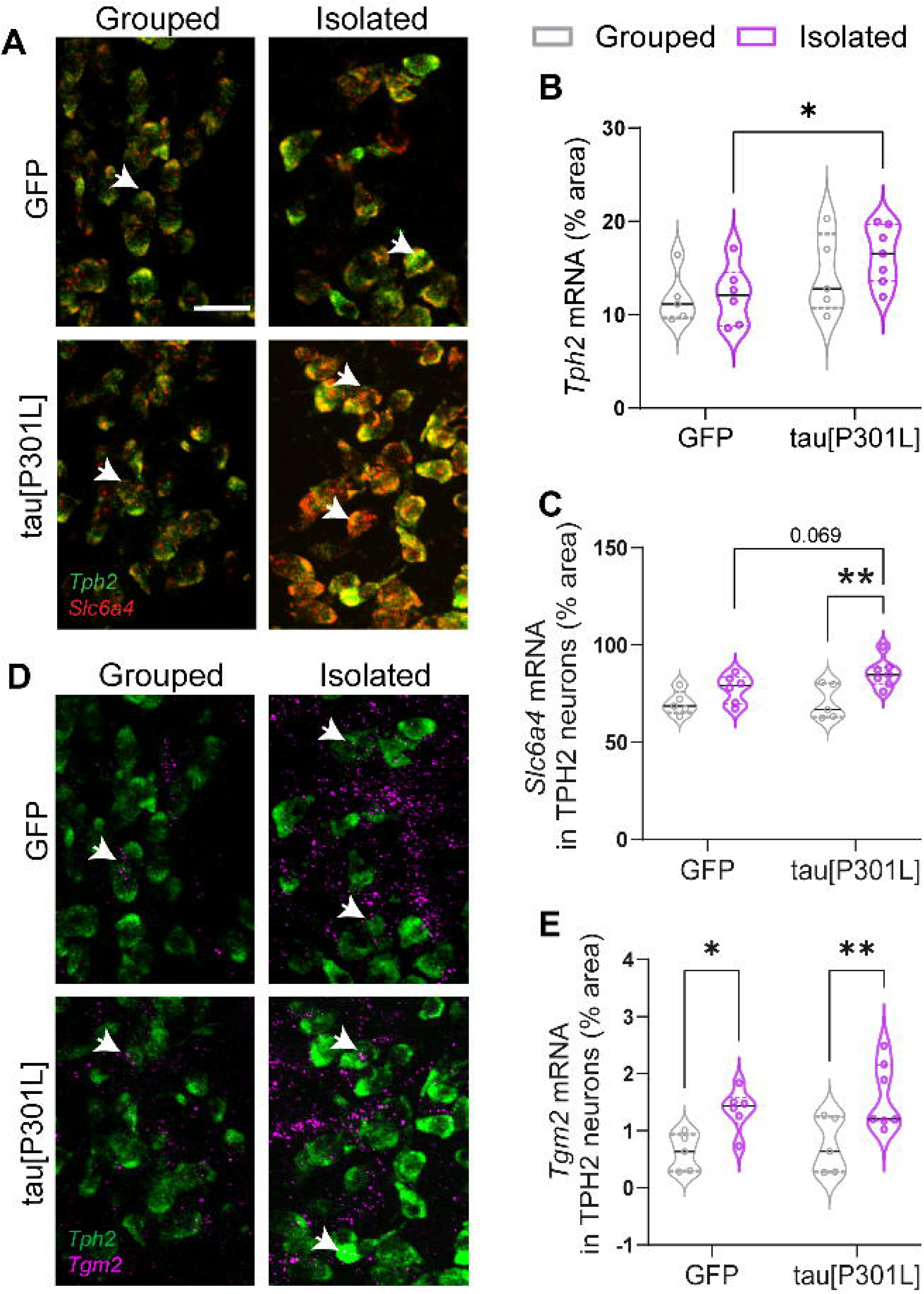
Adolescent isolation stress alters expression of genes related to serotonin synthesis, transport, and stress response in the dorsal raphe nucleus (DRN). (A) Representative images showing *Tph2* and *Slc6a4* mRNA expression in group-housed and socially isolated adolescent GFP and tau[P301L] mice. The top two panels show GFP control: group-housed and isolated conditions, while the bottom two panels show tau[P301L]: group-housed and isolated conditions. Scale bar = 30 µm. Quantification of (B) *Tph2* mRNA levels and (C) *Slc6a4* mRNA levels within TPH2 neurons in grouped and isolated mice injected with GFP or tau[P301L] AAV. (H) Representative images showing *Tgm2* mRNA levels in group-housed and socially isolated adolescent GFP and tau[P301L] mice. The top two panels show GFP control: group-housed and isolated conditions, while the bottom two panels show tau[P301L]: group-housed and isolated conditions. Scale bar = 30 µm. (E) Quantification of *Tgm2* mRNA expression with in TPH2 neurons following adolescent isolation stress. Values (n=7-10/group) are represented as means (±SEM), and the data were analyzed by two-way ANOVA (\**p* < 0.05, \*\**p* < 0.01).

## Adolescent isolation stress drives trans-synaptic spread of tau from DRN to MRN serotonin neurons

We previously reported that C57BL/6J mice injected with AAV-CBA-P301Ltau exhibit increased intrinsic excitability of DRN 5-HT neurons. Heightened neuronal activity, combined with additional stress, may facilitate the trans-synaptic spread of pathological tau through both direct and indirect synaptic connections [16,26–29]. DRN serotonin neurons project extensively to other raphe nuclei, with rostral DRN neurons providing direct inputs to the raphe magnus (RMg) [30,31] and indirect projections to median raphe nucleus (MRN) [32]. We found region-specific alterations in tau pathology and serotonergic neuron density following adolescent social isolation stress. Specifically, stress produced a marked increase in ptau accumulation and TPH2 immunoreactive area in the rostral DRN compared to the mid and caudal subdivisions (Supplementary Figure 1). This led us to further examine tau spread to the MRN and RMg, given the concurrent alterations observed in pain-related behaviors.

The analysis of TPH2 cell density and the percentage of TPH2 immunoreactive area in the MRN revealed no significant group differences across housing or AAV conditions. However, a significantly greater trans-synaptic ptau spread was detected within MRN TPH2 neurons of isolated tau[P301L] mice compared to group-housed tau[P301L] mice (Fig. 4A-E). Two-way ANOVA revealed a significant main effect of virus genotype (*F(*1,28) = 30.79, *p <* 0.0001) and a significant interaction between housing x virus genotype (*F(*1,28) = 5.341, *p =* 0.0284). Post hoc Fisher’s LSD analysis indicated significantly higher ptau levels within TPH2 neurons in isolated tau[P301L] mice compared to group-housed tau[P301L] mice (*p =* 0.0146) (Fig. 4E).

**Figure 4.**
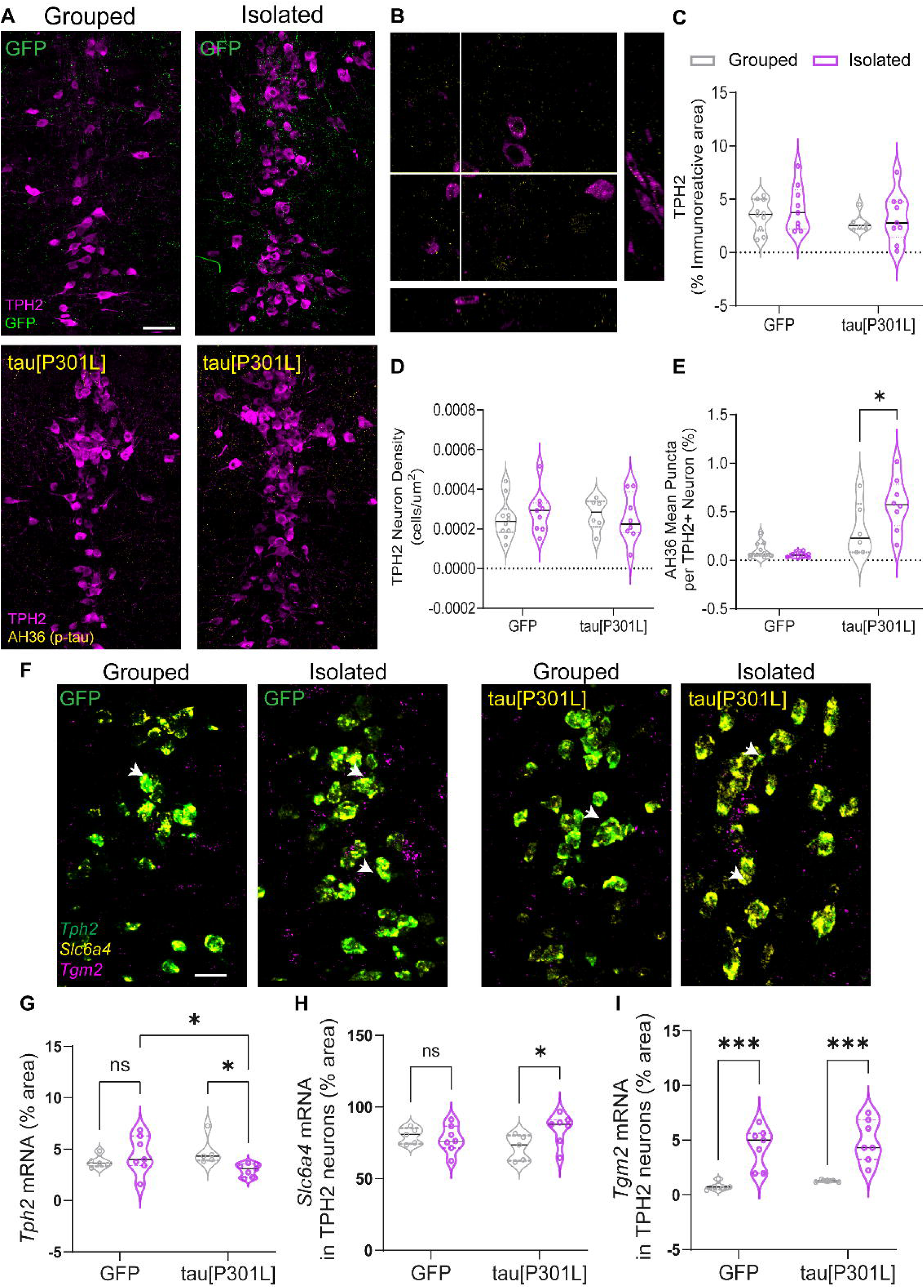
Adolescent isolation stress promotes tau spread from the dorsal raphe nucleus (DRN) to the median raphe nucleus (MRN). (A) Representative confocal images showing increased AH36 (ptau, pSer202/pThr205) immunofluorescence in the MRN of adolescent socially isolated tau[P301L] mice compared to group-housed and GFP controls. The top two panels show GFP control: group-housed and isolated conditions, while the bottom two panels show tau[P301L]: group-housed and isolated conditions. Scale bar = 50 µm. (B) Orthogonal projection image showing ptau accumulation within MRN TPH2-positive neurons. Quantification showing effects of adolescent social isolation on (C) percentage of TPH2-immunoreactive area, (D) TPH2 neuron density, and (E) mean number of AH36 (ptau) puncta per TPH2-positive neuron in GFP and tau[P301L] mice. (F) Representative images showing expression of *Tph2*, *Slc6a4*, and *Tgm2* mRNA in group-housed and isolated adolescent GFP and tau[P301L] mice (left two panels: GFP; right two panels: tau[P301L]). Scale bar = 30 µm. Quantification of *Tph2* (G), *Slc6a4* (H), and *Tgm2* (I) mRNA levels within MRN TPH2-positive neurons in GFP and tau[P301L] mice under group-housed and isolated conditions. Values (n =7-10/group) are represented as means (±SEM), and the data were analyzed by two-way ANOVA (\**p* < 0.05, \*\*\**p* < 0.001).

We next analyzed mRNA levels of *Tph2*, *Slc6a4*, and *Tgm2* in the MRN (Fig. 4F-I). Interestingly, *Tph2* mRNA levels were decreased in isolated tau[P301L] mice compared to both group-housed tau[P301L] and isolated GFP mice. Two-way ANOVA for *Tph2* expression revealed no main effects of virus genotype or housing condition but a significant interaction between housing x virus genotype (*F(*1,20) = 5.642, *p =* 0.0277). Post hoc Fisher’s LSD analysis showed significantly lower *Tph2* mRNA levels in isolated tau[P301L] mice compared to group-housed tau[P301L] (*p =* 0.0242) and isolated GFP mice (*p =* 0.0362) (Fig. 4G). In contrast, *Slc6a4* mRNA expression was selectively altered in tau[P301L] mice. Two-way ANOVA revealed no main effects of virus genotype or housing condition but a trend toward a housing x virus genotype interaction (*F(*1,20) = 3.994, *p =* 0.0595). Post hoc Fisher’s LSD analysis indicated significantly higher *Slc6a4* mRNA levels in isolated tau[P301L] mice compared to group-housed tau[P301L] mice (*p =* 0.0260) (Fig. 4H). Together, these findings suggest that stress and pathological tau synergistically modulate serotonergic gene expression, with reduced *Tph2* and elevated *Slc6a4* expression in isolated tau[P301L] mice indicating disrupted serotonin synthesis and reuptake mechanisms. Finally, we examined *Tgm2* expression in the MRN, given that stress is an established driver of trans-synaptic tau propagation and *Tgm2* is a recognized marker of both stress and AD pathology. Consistent with our findings in the DRN, *Tgm2* mRNA levels were elevated in stressed GFP and tau[P301L] mice. Two-way ANOVA revealed a significant main effect of virus genotype (*F(*1,20) = 35.87, *p <* 0.0001), with no main effect of housing or housing x virus genotype interaction (*F(*1,20) = 0.002842, *p =* 0.9548). Post hoc Fisher’s LSD analysis confirmed significantly higher *Tgm2* mRNA levels in both isolated *tau*[P301L] (*p =* 0.0004) and isolated GFP (*p =* 0.0004) mice compared to their group-housed counterparts (Fig. 4I).

## Adolescent isolation stress drives trans-synaptic spread of tau from DRN to RMg serotonin neurons and its implications in pain behaviors

We next analyzed trans-synaptic tau spread in the RMg serotonergic (TPH2+) neurons. Quantitative analyses of the percentage of TPH2 immunoreactive area, TPH2 cell density, and AH36 (ptau) signal in the RMg revealed alterations in serotonergic markers and tau accumulation in both GFP and tau[P301L] mice. The percentage of TPH2-immunoreactive area was altered across both genotypes, with higher levels in isolated GFP mice and lower levels in isolated tau[P301L] mice (Fig. 5A-E). Two-way ANOVA revealed a significant main effect of housing condition (*F(*1,20) = 8.186, *p =* 0.0097), no main effect of virus genotype, and a trend toward a housing x virus genotype interaction (*F(*1,20) = 3.119, *p =* 0.0926). Post hoc Fisher’s LSD analysis indicated opposing housing effects between isolated GFP mice vs tau[P301L] mice. There was a significantly higher TPH2-immunoreactive area in isolated GFP mice compared to group-housed GFP controls (*p =* 0.0032), and significantly lower TPH2-immunoreactive area in isolated tau[P301L] mice compared to isolated GFP controls (*p =* 0.0098) (Fig. 5C). Similarly, TPH2+ cell density in RMg was influenced by housing conditions in both genotypes. Two-way ANOVA revealed a significant main effect of housing (*F(*1,22) = 4.868, *p =* 0.0381), with no main effect of virus genotype or housing x virus genotype interaction. Post hoc Fisher’s LSD analysis showed a similar opposing pattern between isolated GFP mice vs tau[P301L] mice. There was a significantly higher TPH2 cell density in isolated GFP mice compared to group-housed GFP controls (*p =* 0.0278), and decreasing trend in isolated tau[P301L] mice compared to isolated GFP mice (*p =* 0.0980) (Fig. 5D). Consistent with our findings in the MRN, a significantly greater trans-synaptic ptau spread was detected in RMg TPH2 neurons of isolated tau[P301L] mice compared to group-housed tau[P301L] mice (Fig. 5E). Two-way ANOVA revealed significant main effects of virus genotype (*F(*1,22) = 10.99, *p =* 0.031) and housing condition (*F(*1,22) = 8.559, *p =* 0.0078), as well as a significant interaction between housing x virus genotype (*F(*1,22) = 5.686, *p =* 0.0262). Post hoc Fisher’s LSD analysis confirmed significantly higher ptau levels within RMg TPH2 neurons in isolated tau[P301L] mice compared to group-housed tau[P301L] mice (*p =* 0.0022) (Fig. 5E).

**Figure 5.**
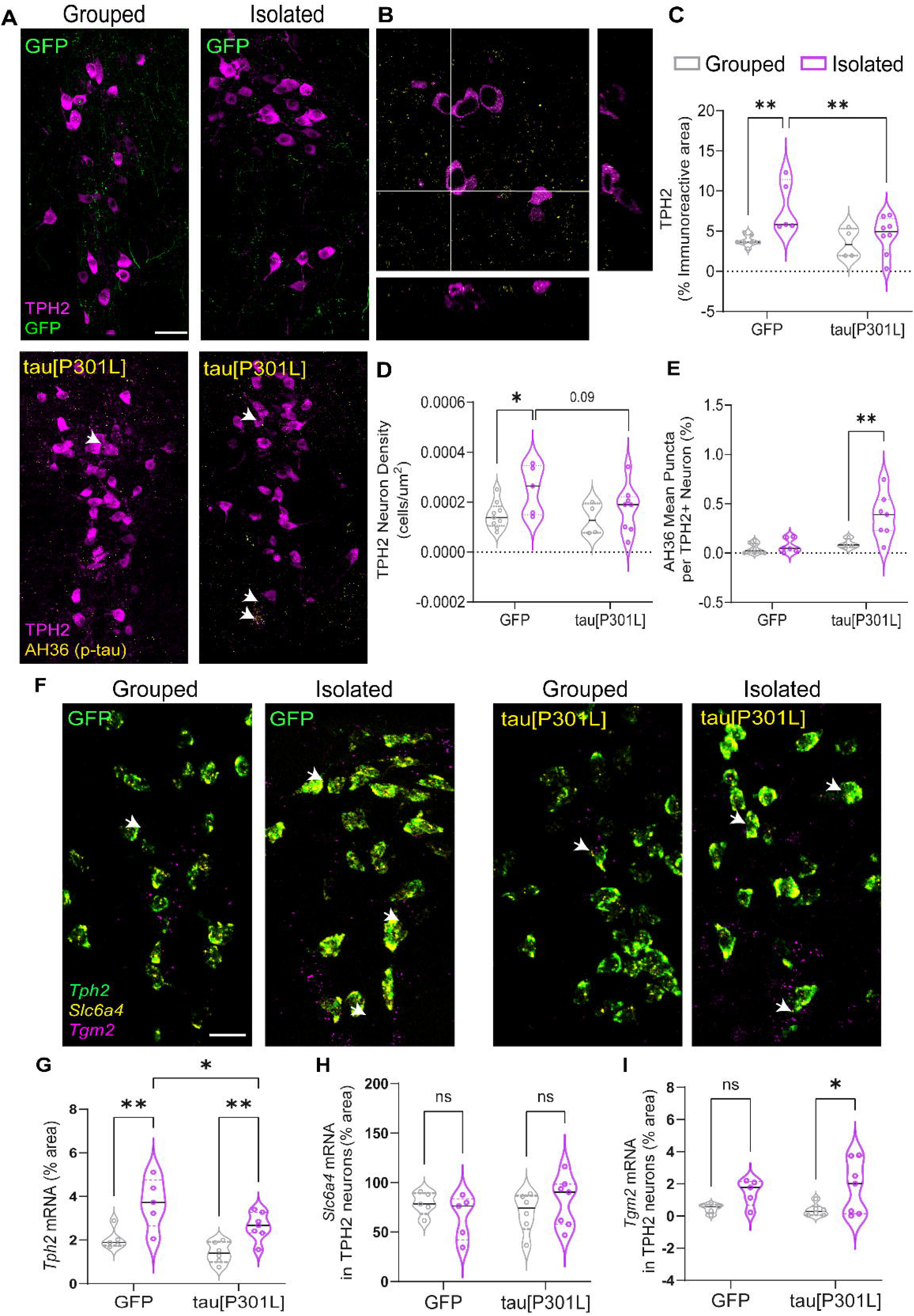
Adolescent isolation stress promotes tau spread from the dorsal raphe nucleus (DRN) to the raphe magnus (RMg). (A) Representative confocal images showing increased AH36 (ptau, pSer202/pThr205) immunofluorescence in the RMg of adolescent socially isolated tau[P301L] mice compared to group-housed and GFP controls. The top two panels show GFP control: group-housed and isolated conditions, while the bottom two panels show tau[P301L]: group-housed and isolated conditions. Scale bar = 50 µm. (B) Orthogonal projection image showing ptau accumulation within RMg TPH2-positive neurons. Quantification showing effects of adolescent social isolation on (C) percentage of TPH2-immunoreactive area, (D) TPH2 neuron density, and (E) mean number of AH36 (ptau) puncta per TPH2-positive neuron within RMg of GFP and tau[P301L] mice. (F) Representative images showing *Tph2*, *Slc6a4*, and *Tgm2* mRNA expression in group-housed and isolated adolescent GFP and tau[P301L] mice (left two panels: GFP; right two panels: tau[P301L]). Scale bar = 30 µm. Quantification of *Tph2* (G), *Slc6a4* (H), and *Tgm2* (I) mRNA levels within RMg TPH2-positive neurons in GFP and tau[P301L] mice under group-housed and isolated conditions. Values (n =7-10/group) are represented as means (±SEM), and the data were analyzed by two-way ANOVA (\**p* < 0.05, \*\**p* < 0.01, \*\*\**p* < 0.001).

We next analyzed mRNA levels of *Tph2*, *Slc6a4*, and *Tgm2* in the RMg (Fig. 5F-I). Interestingly, *Tph2* mRNA levels were increased in both isolated GFP and tau[P301L] mice; however, in isolated tau[P301L] mice there was a significantly lower *Tph2* expression compared to isolated GFP mice, similar to MRN findings. Two-way ANOVA for *Tph2* expression revealed significant main effects of virus genotype (*F(*1,19) = 7.891, *p =* 0.0112) and housing condition (*F(*1,19) = 22.50, *p =* 0.0001), with no significant interaction between housing x virus genotype. Post hoc Fisher’s LSD analysis indicated significantly higher *Tph2* mRNA levels in both isolated GFP (*p =* 0.0018) and tau[P301L] (*p =* 0.0064) mice compared to their respective group-housed counterparts, as well as a significant reduction in isolated tau[P301L] compared to isolated GFP mice (*p =* 0.0207) (Fig. 5G). In contrast, *Slc6a4* mRNA expression was unaltered across all groups. Two-way ANOVA showed no significant main effects of virus genotype, housing condition, or housing x virus genotype interaction (Fig. 5H). We next assessed *Tgm2* expression in the RMg. Unlike in the DRN and MRN, where *Tgm2* expression was elevated in both isolated GFP and tau[P301L] mice, *Tgm2* mRNA levels in the RMg were significantly increased only in isolated tau[P301L] mice. Two-way ANOVA revealed a significant main effect of housing condition (*F(*1,19) = 7.871, *p =* 0.013), with no effects of virus genotype or interaction between housing x virus genotype. Post hoc Fisher’s LSD analysis confirmed significantly higher *Tgm2* mRNA levels in isolated tau[P301L] mice compared to group-housed tau[P301L] mice (*p =* 0.0226) (Fig. 5I).

Together, these findings suggest that stress and pathological tau synergistically modulate serotonergic gene expression in the RMg. The reduction in TPH2 cell density and the altered *Tph2* expression pattern in isolated tau[P301L] mice, relative to GFP controls, indicate potential serotonergic neuronal loss as a consequence of trans-synaptic tau spread mechanisms that may underlie increased stress and pain sensitivity observed in behavioral outcomes.

## Discussion

This study identifies a mechanistic pathway through which adolescent psychosocial stress contributes to early neuropsychiatric and sensory alterations in AD. Social withdrawal during adolescence represents a major environmental factor within the exposome that can rewire serotonergic circuits involved in mood, depression, and pain regulation. Using an AAV-based approach to express tau[P301L] in the DRN, we examined the impact of adolescent psychosocial stress on tau accumulation and its trans-synaptic spread to other raphe nuclei. This combination of stress and tau pathology produced robust neuropsychiatric impairments and pain sensitivity. These findings position the DRN as a critical, stress-sensitive hub through which environmental exposures such as social isolation or loneliness shape long-term vulnerability to AD-related dysfunction.

Our results are built on postmortem and preclinical evidence demonstrating that tau pathology often initiates in the DRN, an area involved in mood and pain regulation, well before the onset of cognitive symptoms [3,7,9,13,33]. Given the serotonergic system’s established role in affective and pain disorders, the present findings extend this framework by demonstrating that adolescent social stress enhances tau propagation from the DRN to interconnected serotonergic nuclei, including the MRN and RMg. These regions regulate social behavior, anxiety, and descending pain pathways; thus, their dysfunction likely contributes to the constellation of neuropsychiatric and sensory abnormalities observed in prodromal AD [4,6,34,35]. Our prior work demonstrated that DRN tau[P301L] transduction induces social deficits, while adolescent isolation independently produces anxiety- and depression-like behaviors [9,20]. Consistent with these findings, the present study shows that adolescent isolation alone increased anxiety-like behavior, while social impairments were observed only in tau[P301L] mice and were further exacerbated by adolescent isolation. These data indicate that raphe tau pathology, in combination with social stress, synergistically drives social dysfunction. Moreover, both GFP and tau[P301L] groups exhibited hyperlocomotion in the open field test, consistent with previous studies [9,20,36]. These findings reinforce that serotonergic circuits are particularly sensitive to adolescent stress, conferring susceptibility to mood disorders and facilitating tau spread into downstream raphe nuclei enriched in serotonergic neurons [36–38].

We next examined whether stress influences tau phosphorylation within 5-HT neurons and how this affects 5-HT activity. Our previous work demonstrated that tau[P301L] transduction in the DRN negatively correlates with TPH2 cell density and ptau accumulation, although overall TPH2 levels remained unchanged in stress-naïve mice. In the current study, however, stress elevated TPH2 expression and enhanced ptau accumulation within 5-HT neurons. This aligns with our earlier observation that tau[P301L]-transduced DRN neurons are hyperexcitable under stress-naïve conditions [9]. Together, these findings suggest that under stress, tau-transduced 5-HT neurons not only exhibit hyperexcitability but also increase serotonin reuptake, as indicated by elevated *Slc6a4* (SERT) expression within TPH2-positive neurons in socially isolated tau groups. This pattern likely reflects a compensatory response to reduced extracellular serotonin levels under chronic stress. Such compensatory upregulation of serotonergic signaling has been reported in rodent models of anxiety and major depressive disorder, where increased *Tph2* mRNA and protein expression were observed in Sprague-Dawley rats displaying social deficits following inescapable tail-shock stress [39]. Similarly, chronic social defeat stress has been shown to increase TPH2 cell density in the DRN of adult male rats exhibiting anhedonic behavior [40]. These findings support the notion that adolescent social isolation can act as a potent environmental stressor that induces depressive-like phenotypes before the onset of cognitive impairment in AD. The concurrent increase in *Tgm2*, an established stress and tau aggregation gene in stressed GFP and tau[P301L] mice, further supports a mechanism whereby stress amplifies tau aggregation and neuronal vulnerability in serotonergic circuits, as reported by our group in a *Htau* mouse model [13]. The serotonergic system thus emerges as a particularly vulnerable target during the prodromal stages of AD, where stress-induced dysregulation may promote tau accumulation and trans-synaptic spread to downstream circuits.

DRN projects extensively to other raphe nuclei, with rostral neurons sending direct inputs to the RMg [30,31] and indirect projections to MRN [32]. Some studies also suggest that caudal DRN neurons share molecular and functional similarities with MRN neurons [41]. In the present study, we observed stress-specific effects on tau pathology and TPH2 density predominantly within the rostral DR, which exhibited a higher density of TPH2 positive neurons and elevated ptau levels compared to the mid or caudal regions. Interestingly, similar stress-related increases in ptau accumulation and altered TPH2 neuron density were also evident in the caudal DR, prompting us to examine projections and the trans-synaptic tau spread to MRN and RMg, given the concurrent alterations observed in pain-related behaviors (Supplementary Fig. 1).

Within the MRN, *Tph2* mRNA levels were markedly reduced in adolescent isolated tau[P301L] mice compared to GFP and group-housed controls, whereas *Slc6a4* (SERT) mRNA levels were elevated. This pattern suggests a compensatory mechanism maintaining steady TPH2 protein levels within serotonergic neurons despite transcriptional dysregulation in the isolated tau[P301L] mice. Additionally, stress exposure increased *Tgm2* mRNA expression in both GFP and tau[P301L] mice, consistent with its established role as a molecular indicator of stress [22] and as a crosslinking enzyme that promotes phosphorylated tau aggregation [42–44]. Stress-induced *Tgm2* upregulation may therefore exacerbate ptau accumulation, neuronal loss, and disease progression, contributing to anxiety-related phenotypes in conjunction with DRN serotonergic dysfunction. This interpretation aligns with prior findings showing that activation or inhibition of serotonergic neurons in the DRN and MRN, respectively, are associated with depressive-like behaviors in rodent models [45].

The main function of RMg is to modulate pain by sending projections to the dorsal horn of the spinal cord [46]. A major finding of this study is that ptau propagates trans-synaptically from the DRN to the RMg, altering serotonergic modulation and producing changes in thermal and mechanical pain sensitivity. This underscores a critical interaction between stress, serotonergic dysfunction, and raphe tau pathology in shaping pain outcomes in AD. Adolescent socially isolated tau[P301L] mice exhibited increased thermal and mechanical sensitivity in the Hargreaves and von Frey tests compared with isolated GFP controls. This is noteworthy given that pain is often under-recognized in early AD, as affected individuals are less likely to self-report pain due to impaired pain-memory associations [4,34,47]. To our knowledge, this is the first preclinical evidence linking trans-synaptic tau spread to selective alterations in mechanical pain sensitivity, paralleling the reduced pain awareness commonly reported in early AD.

The distinct effects of stress alone versus combined stress and tau pathology on nociception suggest that different serotonergic and spinal pathways are disrupted when these factors co-occur. Serotonergic projections from the DRN to the RMg shape both spinal and supraspinal pain processing, and our data indicates that tau propagation within these circuits interferes with this regulation. Stress alone increased TPH2 density in the RMg, whereas stressed tau[P301L] mice showed a marked loss of TPH2-positive neurons accompanied by ptau accumulation.

These results support a model in which DRN-derived tau spreads to the RMg, reducing serotonergic output and initiating neurodegenerative signaling that weakens nociceptive transmission in the dorsal horn. Complementary work from our group shows that adolescent alcohol exposure similarly diminishes RMg 5-HT density and heightens pain sensitivity [48], highlighting the vulnerability of this system to environmental stressors. In the present study, however, nociceptive deficits emerged only in stressed tau[P301L] mice, suggesting that serotonergic loss is the key point of convergence through which stress and tau pathology impair downstream pain circuits. Other studies have similarly reported that the loss of 5HT in RMg is linked to hyperalgesia effect in other pain models (diabetic-induced neuropathy and sciatic nerve ligation) [49]. Together, these findings demonstrate that stress-induced hyperexcitability and tau-dependent neurodegeneration within raphe circuits act synergistically to alter serotonergic control of pain, producing modality-specific nociceptive changes reminiscent of early sensory deficits in AD. Future studies should assess how tau pathology influences RMg 5-HT function and dorsal horn 5-HT receptor engagement during stress-induced hyperalgesia in tau/AD models.

## Conclusion

This study demonstrates that adolescent psychosocial stress, a key exposomic factor, amplifies tau propagation within brainstem serotonergic circuits, producing early neuropsychiatric and sensory changes relevant to AD. Stress-induced tau accumulation in the DRN and its projections to the MRN and RMg led to anxiety-like behavior, social deficits, and disrupted pain regulation. These results identify the DR → MRN and RMg serotonergic axis as a stress-sensitive hub where environmental exposure and tau pathology converge to drive mood and pain dysfunction. Together, these findings provide a mechanistic framework linking adolescent stress, tau spread, and early vulnerability to Alzheimer’s-related neuropsychiatric and pain disturbances.

## Declarations

Ethics approval: All procedures on mice in this study were approved by the Institutional Care and Use Committee at the University of Iowa and University of Florida.

Availability of data and materials: All data generated or analyzed during this study are included in this published article and its supplementary information files.

## CRediT authorship contribution statement

**C.A.M:** Conceptualization, Writing – review & editing, Visualization, Software, Methodology, Investigation, Formal analysis, Data curation, Funding acquisition, Supervision. **N.B:** Writing – Conceptualization, Writing - original draft, review & editing, Visualization, Software, Methodology, Investigation, Formal analysis, Data curation, Supervision. **G.G.:** Writing – parts of original draft, Visualization, Software, Methodology, Investigation, Formal analysis, Data curation. **K.K:** Methodology, Investigation, Formal analysis, Data curation. **N.B (Biggerstaff):** Methodology, Investigation, Formal analysis, Data curation.

