## Supplementary figure 1 for "Adolescent Social Isolation Facilitates Tau Spread in Raphe Nuclei, Linking Depression and Hyperalgesia in Alzheimer’s Disease"

### Supplementary Files

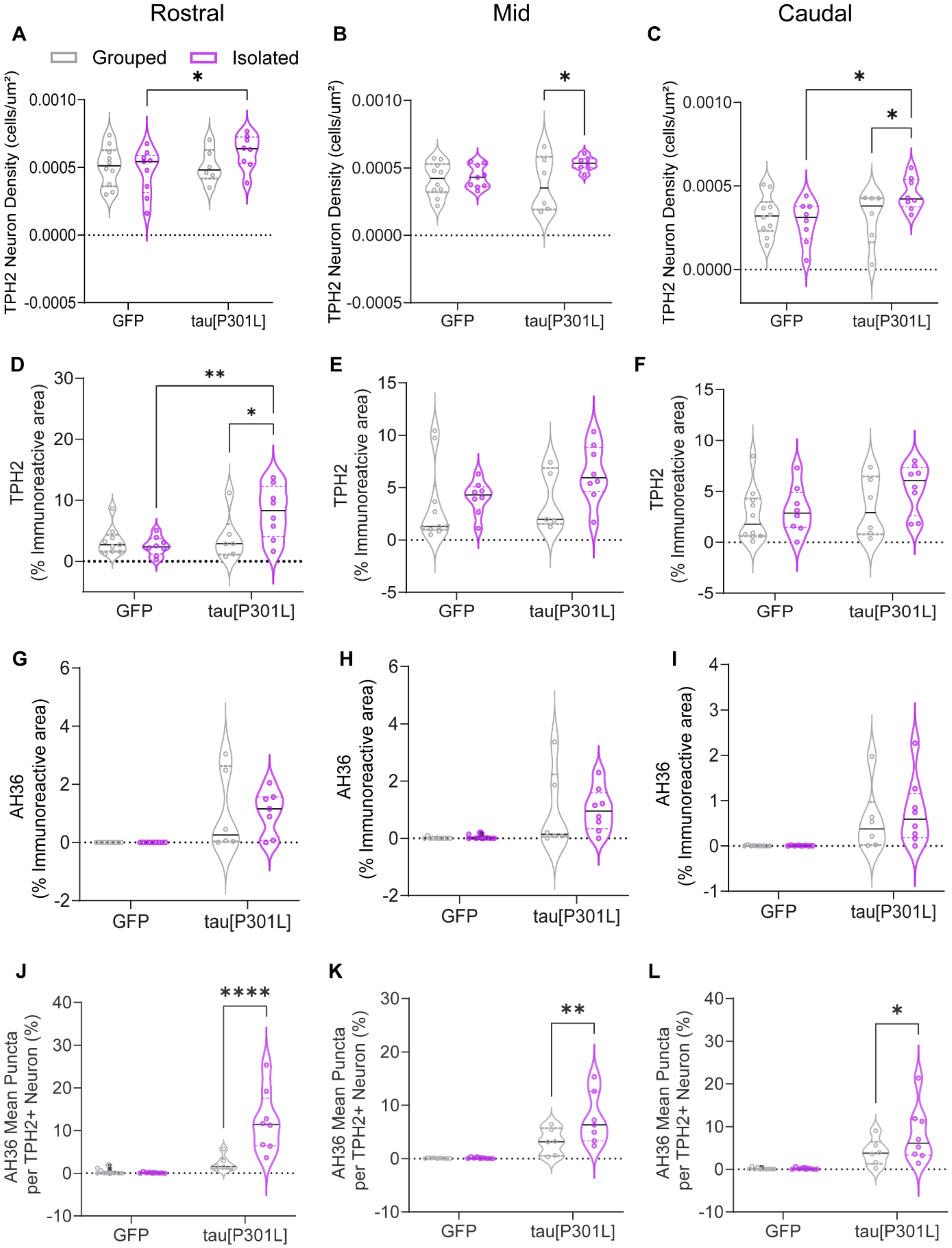

**Supplementary Fig. 1:** Quantification showing the effects of adolescent social isolation on rostral to caudal dorsal raphe nucleus (DRN) axis on (A-C) TPH2 neuron density, (D-F) TPH2-immunoreactive area, (G-I) percentage of AH36-immunoreactive area, and (J-K) mean number of AT8 (ptau) puncta per TPH2-positive neuron in GFP and tau[P301L] mice. Values (n =7-10/group) are represented as means (±SEM), and the data were analyzed by two-way ANOVA (*$p < 0.05$, **$p < 0.01$, ****$p < 0.0001$).
